# Structure and mechanism of a Type III CRISPR defence DNA nuclease activated by cyclic oligoadenylate

**DOI:** 10.1101/784280

**Authors:** Stephen A McMahon, Wenlong Zhu, Shirley Graham, Robert Rambo, Malcolm F White, Tracey M Gloster

## Abstract

The CRISPR system provides adaptive immunity against mobile genetic elements in prokaryotes. On binding invading RNA species, Type III CRISPR systems generate cyclic oligoadenylate (cOA) molecules which act as a second messenger, signalling infection and potentiating a powerful immune response by activating a range of downstream effector proteins that can lead to viral clearance, cell dormancy or death. Only one type of effector enzyme has been studied – the Csm6/Csx1 ribonuclease domain, and the mechanism of cOA activation is not understood at a molecular level. Here we describe the structure and mechanism of a novel cOA-activated CRISPR defence DNA endonuclease, Can1 (“CRISPR ancillary nuclease 1”). Can1 has a unique monomeric structure with two CRISPR associated Rossman fold (CARF) CARF domains and two DNA nuclease-like domains. The crystal structure of the enzyme has been captured in the activated state, with a cyclic tetra-adenylate (cA_4_) molecule bound at the core of the protein. cA_4_ binding reorganises the structure to license a metal-dependent DNA nuclease activity specific for nicking of supercoiled DNA. DNA nicking by Can1 is predicted to slow down viral replication kinetics by leading to the collapse of DNA replication forks.

## INTRODUCTION

Type III (Csm/Cmr) CRISPR systems utilise a cyclase domain in the Cas10 subunit to generate cyclic oligoadenylate (cOA) by polymerising ATP^1–3^. The cyclase activity is activated by target RNA binding and switched off by subsequent RNA cleavage and dissociation^1,4^. cOA is a potent anti-viral second messenger, that sculpts the anti-viral response by binding to and activating CRISPR associated Rossman fold (CARF) family proteins^5^. The only CARF family enzymes to be characterised to date are dimeric ribonucleases of the Csx1/Csm6 family, which have a C-terminal HEPN (Higher Eukaryotes and Prokaryotes, Nucleotide binding) domain that is allosterically activated by cOA binding in the CARF domain^6–9^. Csm6 from *Enterococcus italicus* and *Streptococcus thermophilus* are activated by cyclic hexa-adenylate (cA_6_)^2,3^, whilst Csx1 from *Sulfolobus solfataricus* and Csm6 from *Thermus thermophilus* are activated by the cyclic tetra-adenylate (cA_4_) molecule^1,2,10^. In addition to ribonucleases, a wide range of CARF family proteins have been identified, including transcription factors such as Csa3^11^, cA_4_-degrading ring nucleases^12^ and putative membrane proteins of unknown function^5,13^, suggesting that much remains to be discovered. The cOA signalling pathway is crucial for defence against mobile genetic elements (viruses and plasmids) in multiple different systems^7,14–17^. In infected cells, activation of Csm6 results in degradation of both host and invading RNA, leading to significant reductions in cell growth until infection is cleared^17^.

*T. thermophilus* has well characterised type III CRISPR systems and accessory proteins. The *T. thermophilus* Cmr (type III-B) complex was one of the first studied biochemically, revealing the distinctive cleavage of target RNA with 6 nucleotide spacing^18,19^. High resolution cryo-EM studies revealed the mechanistic basis of this RNA cleavage by the backbone subunit Cmr4^20^. The *T. thermophilus* Csm (Type III-A) complex was shown to possess both sequence specific RNA cleavage and RNA-dependent DNA degradation activities, fitting the emerging consensus for type III CRISPR defence^21^. The crystal structure of the *T. thermophilus* Csm6 (TTHB152) protein revealed a dimeric arrangement with a HEPN nuclease domain possessing weak ribonuclease activity and a CARF domain postulated to bind activating ligands^6^. This was subsequently confirmed with the discovery that the cyclase domain of Cas10 generates cyclic oligoadenylates that activate CARF domain proteins, including Csm6, which is stimulated by cA_4_ binding to become a much more active ribonuclease^2^. A second Csm6 orthologue (TTHB144) was recently shown to be activated by cA_4_ binding and also capable of cA_4_ degradation – the first example of a self-limiting Csm6-family protein^10^.

TTHB155, hereafter Can1 (CRISPR ancillary nuclease 1), was identified as a potential accessory protein of the type III CRISPR system by Shah *et al.* in 2018 (cluster 107)^22^. The gene is strongly up-regulated following phage infection in *T. thermophilus* and is situated adjacent to the CRISPR-5 locus, with operons encoding subunits of the Cmr and Csm complexes nearby^23^. Can1 is a large protein, 636 amino acids in length. Sequence analysis using HHPRED^24^ suggested the presence of two CARF domains, one at the N-terminus and one towards the centre of the protein, together with a C-terminal PD-D/ExK family metal-dependent nuclease domain (Figure 1A). The protein therefore has a number of unique features that have not been investigated previously. Here, we show that Can1 is a metal dependent DNA nuclease, activated by cA_4_ binding. The structure of Can1 bound to cA_4_ reveals extensive structural reorganisation around the activator, bringing the two CARF domains together and forming a DNA binding interface with an active site for DNA cleavage. The enzyme preferentially nicks supercoiled DNA to generate ligatable products, which may slow down viral replication without causing catastrophic damage to the host genome.

**Figure 1.**
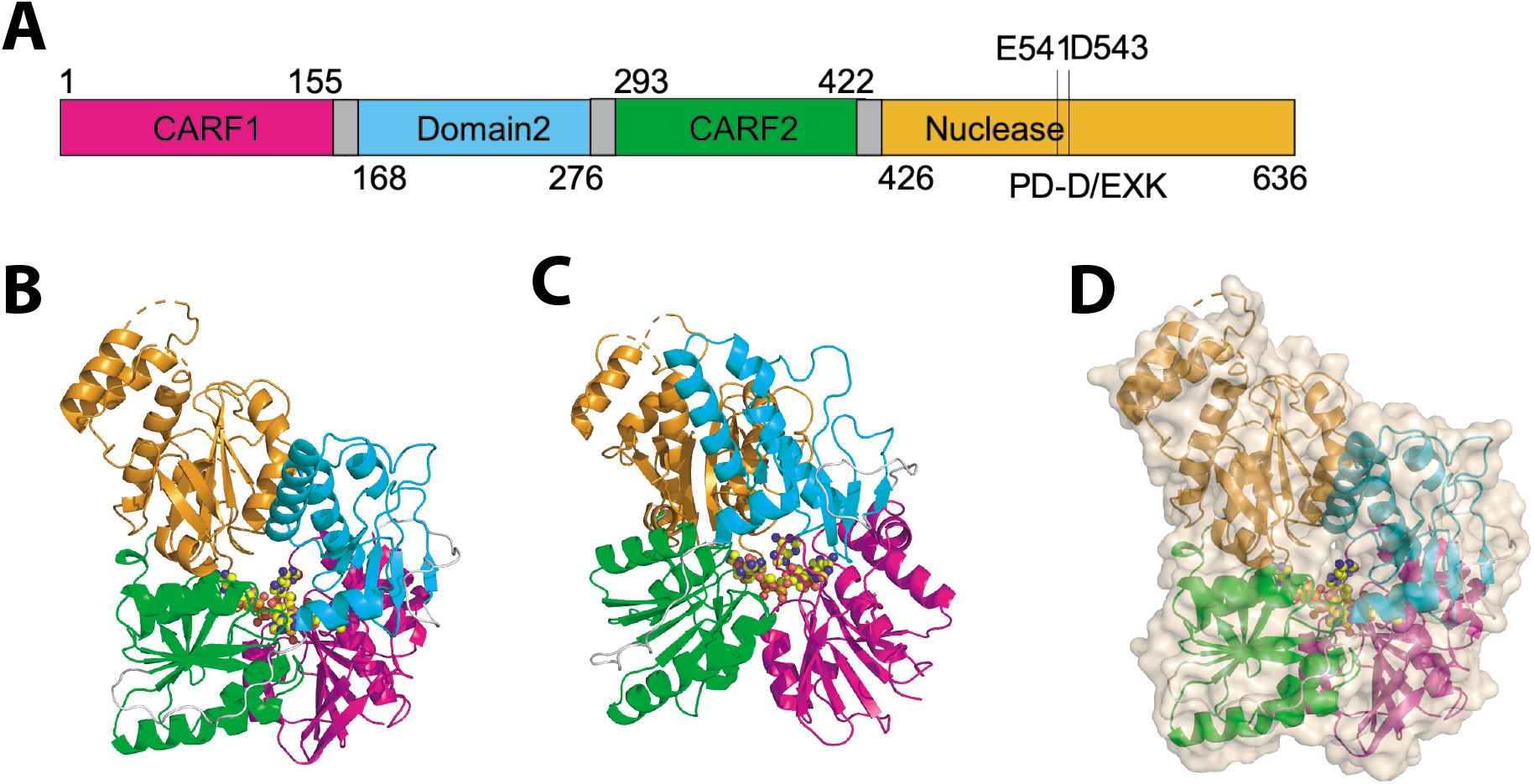
Domain organisation and structure of Can1. **(A)** Domain organisation of Can1 with residue numbers indicated. **(B)** and **(C)** Overall structure of Can1 shown in cartoon representation. Coloured as in panel (A): first CARF domain (magenta), domain 2 (cyan), second CARF domain (green), nuclease domain (orange), loops between domains (grey). A molecule of cA_4_ bound to Can1 is shown in yellow sphere representation. **(D)** Overall structure of Can1, as coloured in panel (B), with surface representation to illustrate cA_4_ is completely buried.

## RESULTS

### Overall structure of Can1 bound to cA_4_

A synthetic gene encoding Can1 was constructed and cloned into the *E. coli* expression vector pEV5HisTEV^25^. The protein was expressed with an N-terminal, TEV protease-cleavable polyhistidine tag, and purified by immobilised metal affinity chromatography and gel filtration, with concomitant tag removal, as described in the Methods. Given the observation that the *T. thermophilus* Csm6 ribonucleases are activated by cA_4_^2,10^, we reasoned that Can1 might bind the same activator. Can1, with selenomethionine incorporated, was crystallised in the presence of cA_4_. Diffraction data were collected at Diamond Light Source to 1.83 Å resolution and phased using the incorporated selenium atoms. Following autobuilding of the model, it was refined to a final R_work_ of 17.7% and R_free_ of 21.1%. The protein chain is traced from residue 3 to 638, with three short sections too disordered to model (464-471, 491-494 and 530-539). Efforts were made to crystallise apo Can1, but no crystals were forthcoming.

The structure of Can1 is unique, with four distinct domains linked by flexible loops (Figure 1). At the N-terminus there is a CARF domain (residues 3-151), domain 2 (residues 168-276), a second CARF domain (residues 293-422) and a nuclease domain (residues 426-638). The two CARF domains display the canonical Rossman fold, each with a core of five parallel β-strands alternating with four α-helices. This is followed by a further two β-strands that form a beta-hairpin, and an α-helix. The nuclease domain at the C-terminus comprises a central core of six β-strands flanked by six α-helices typical of the PD-D/ExK metal-dependent nuclease family^26^, with an additional three helices on the periphery. Domain 2 comprises five β-strands flanked by four α-helices, reminiscent of part of the core domain of the PD-D/ExK nuclease family, with a helix-turn-helix motif between the second and third helices.

### cA_4_ recognition by Can1

There is unambiguous electron density in the *F*_obs_ – *F*_calc_ map at 3σ corresponding to a molecule of cA_4_ bound to Can1 (Figure 2A), which was modelled following generation of the library in Acedrg^27^. As this is the first example of a cOA molecule visualised at atomic resolution, we present a detailed investigation of the interactions observed. The molecule of cA_4_ sits at the interface between the two CARF domains of Can1 and is completely enclosed by the protein (Figure 1C). There are a fairly modest number of hydrogen bond interactions between Can1 and cA_4_ given its size, all but one of which originate from the two CARF domains (Figure 2B and Figure S1). There are hydrogen bonds formed between three of the AMP molecules in cA_4_ and residues in the first CARF domain; these interactions are formed via main chain atoms of residues Lys90, Asn12, Asp13 and Gly88, and side chain atoms of residues Lys90, Asn12, Tyr143, His113 and Tyr147. Likewise, there are hydrogen bond interactions formed between three of the AMP molecules in cA_4_ and residues in the second CARF domain; these are formed via main chain atoms of residues Thr384, Gln301, Gly405 and Asn380, and side chain atoms of residues Tyr402, Thr384, Glu324, Thr322, Ser299 and Asn380. A total of 19 water molecules also hydrogen bond directly with cA_4_.

**Figure 2.**
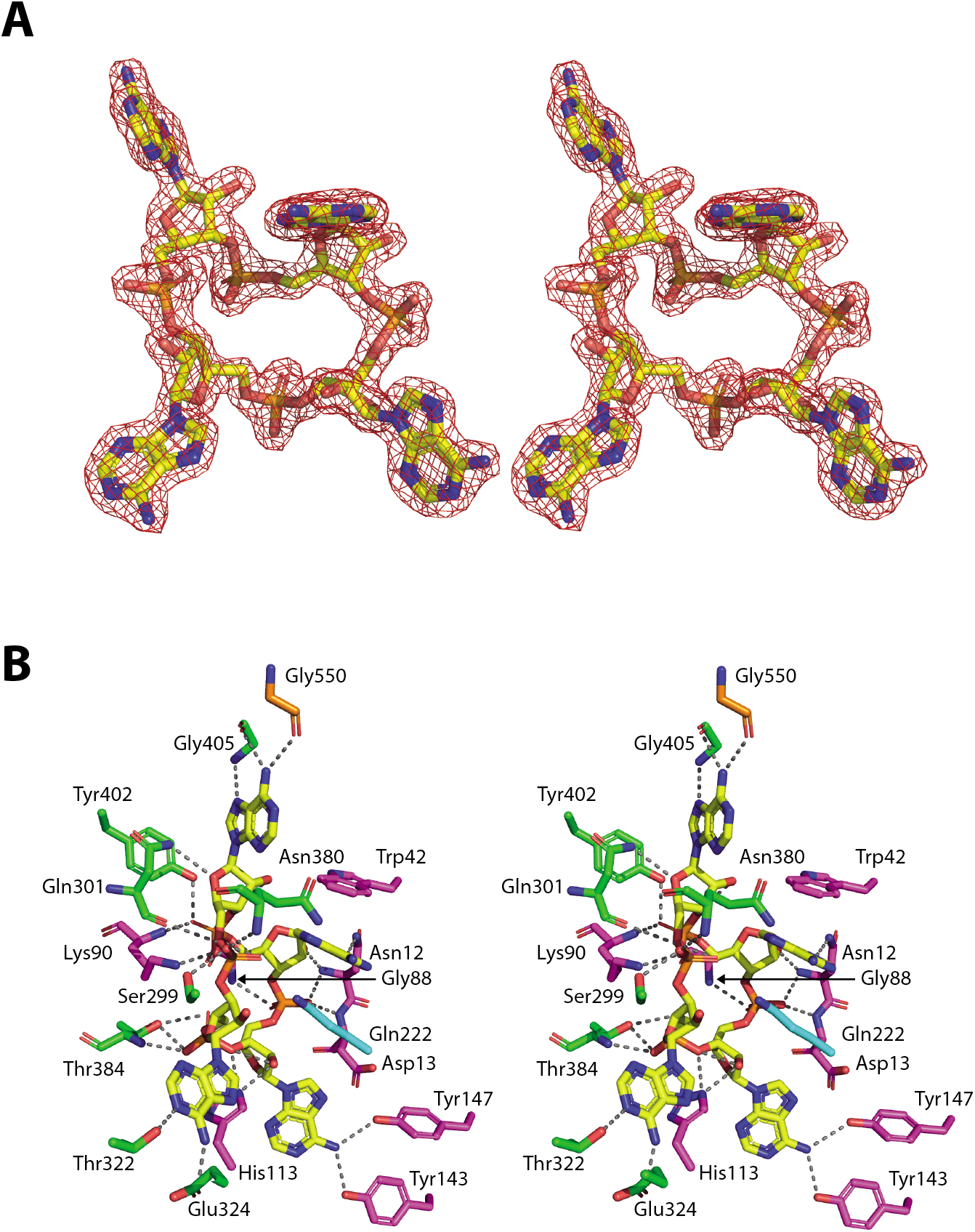
Structure of the cA_4_ activator and interactions with Can1. **(A)** Divergent stereo representation of cA_4_ (in yellow stick representation) with the maximum likelihood/*σ_A_* weighted 2*F*_obs_ – *F*_calc_ electron density map contoured at 1.7 sigma shown in red. **(B)** Divergent stereo representation of the binding site of cA_4_ in complex with Can1. cA_4_ and binding site residues are shown in stick representation, with cA_4_ in yellow and amino acid residues coloured by the domain from which they originate (CARF domain 1: magenta, domain 2: cyan, CARF domain 2: green, nuclease domain: orange). The dotted lines represent hydrogen bond interactions.

Superposition of the four adenosine units (centred on the carbon atoms in the ribose sugar) within cA_4_ show a marked difference in the conformation of one of them (Figure S2). Whilst three of the adenosine molecules superimpose well (with respect to the ribose sugar and imidazole moiety of the adenine base; one adenine base is flipped compared to the other two but in the same plane), the adenine of the fourth adenosine molecule is in a more axial position at the anomeric carbon. This adenine base displays a clear π-π stacking interaction with Trp42, which appears to be the driving force for this alternate, and most probably higher energy, conformation. It has the knock-on effect of causing the ribose sugar to display a 2’-exo conformation, compared to 2’-endo conformation, which is observed in the other adenosine units.

Only two residues from the nuclease domain and domain 2 are in close proximity to cA_4_. Gly550, from the nuclease domain, forms a hydrogen bond interaction via its main chain carboxyl group to an adenine base (Figure 2B). Gln222 from domain 2 does not interact directly with any atoms in cA_4_, but hovers above the centre of the cA_4_ molecule, and makes two water-mediated interactions to opposite phosphate groups in cA_4_ (Figure S3).

All other characterised CARF family proteins consist of a single CARF domain, which homo-dimerizes to bind one molecule of cOA. Can1 is unique as it possesses two CARF domains in a single polypeptide. The CARF domains superimpose with an RMSD of 2.5 Å over 113 amino acids, although few residues are conserved (Figure S4, S5A). A striking feature of cA_4_ recognition is the asymmetry of the interactions formed with Can1, as the interactions with each AMP molecule are distinct. The comparison highlights the structural equivalence of Asp13 and Gln301, which both make main chain interactions with cA_4_, and of Lys90 and Thr384, which both make main chain and side chain interactions with cA_4_.

### Structural comparisons suggest the cA_4_ binding site is pre-formed in the apo protein

DALI^28^ searches for the nearest structural homologue to each CARF domain in Can1 gave identical results for each: hypothetical protein VC1899 (PDB: 1XMX), a CARF family protein that is a member of the DUF1887 family. The CARF dimer (taken as the two domains from a single polypeptide chain) of Can1 superimposes with the CARF dimer (formed by two monomers) of VC1899 with an RMSD of 3.6 Å over 294 residues (Figure 3A). This superimposition allows exploration of likely interactions between VC1899 and cA_4_. All of the main chain interactions observed between Can1 and cA_4_ are likely conserved, as are a number of the side chain interactions (Figure S4 and S5B). There is no equivalent in VC1899 to Trp42, the residue responsible for causing one of the adenine bases to sit in a more axial conformation with respect to the ribose sugar. With the absence of this interaction this adenine may not be in such a strained position when bound to VC1899, and there is sufficient room in the binding site for this to fit in a more relaxed position (Figure 3B). Few close interactions/clashes between cA_4_ and VC1899 are predicted, and these involve side chains which could easily be repositioned, suggesting there is little rearrangement of either the CARF dimer or of residues in the CARF domains upon binding of cA_4_. In other words, the cA_4_ binding site is likely to be largely pre-formed in the apo-protein.

**Figure 3.**
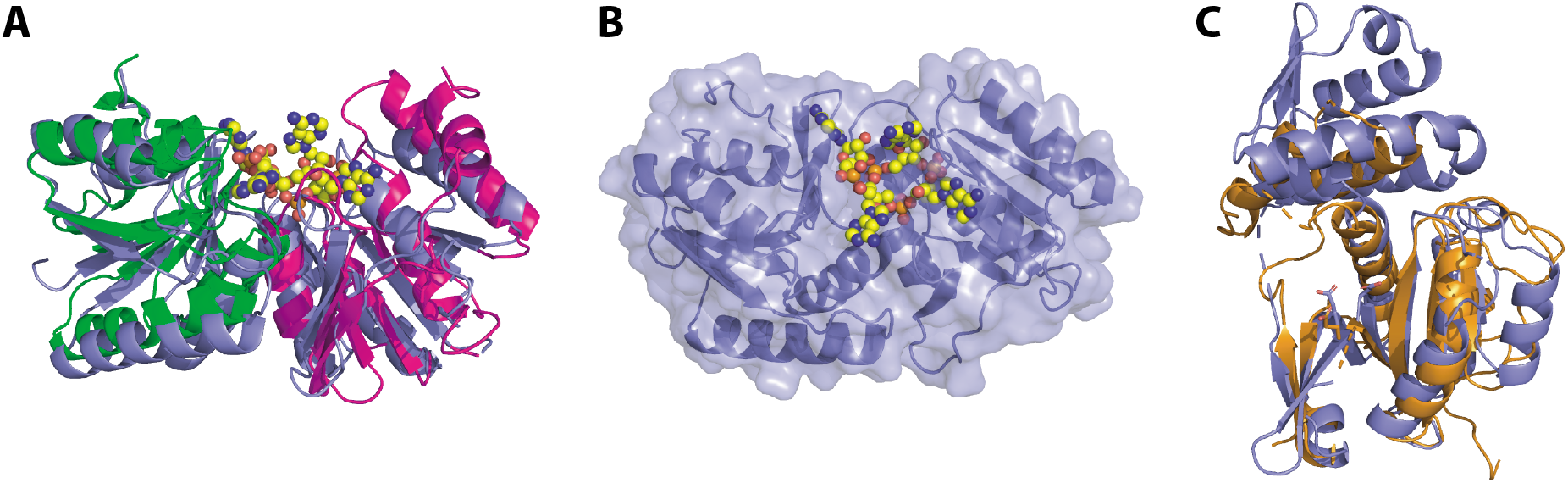
Structural comparison of Can1 and the DUF1887 family protein VC1899. **(A)** Superimposition of the CARF domains from Can1 (coloured as in Figure 1) in complex with cA_4_ (yellow spheres) with the CARF domain from VC1899 (mauve). The VC1899 dimer (monomer plus the symmetry mate that forms the functional dimer) was overlapped with Can1. **(B)** Surface and cartoon representation of VC1899, with cA_4_ (yellow spheres) modelled based on the superimposition shown in panel (A). **(C)** Superimposition of the nuclease domain from Can1 (orange) and the nuclease domain from VC1899 (mauve). The active site residues are shown in sticks.

### The Nuclease and nuclease-like domains

DALI searches for structural homologues of the nuclease domain of Can1 once again gave the hypothetical protein VC1899 (PBD: 1XMX) as the highest hit. The nuclease domain from Can1 superimposes with the nuclease domain of VC1899 with an RMSD of 3.4 Å over 160 residues (Figure 3C, with nuclease active site residues shown). Whilst the core of the nuclease domain is very similar, Can1 has an extra helix-loop-helix feature between residues 441 and 498, whereas at the equivalent position (residues 156-248) VC1899 possesses a non-superimposable section. The active site residues Glu541, Asp543, Glu560 and Lys562 in Can1 overlap with equivalent active site residues in VC1899. The closest structural homologues with dsDNA bound are the type 2 restriction enzymes AgeI (PDB: 5DWB^29^) and NgoMIV (PDB: 1FIU^30^), which overlap with the nuclease domain of Can1 with an RMSD of 4.0 Å over 143 residues and 3.4 Å over 133 residues, respectively (Figure S6). The core secondary structure elements of the nuclease domain of Can1 and each of the restriction enzymes are conserved, and the active site residue Asp543 in Can1 overlaps with Asp142 in the AgeI nuclease and Asp140 in the NgoMIV nuclease. Thus, the approximate binding position of the dsDNA in complex with Can1 can be predicted with some confidence (Figure 4A).

**Figure 4.**
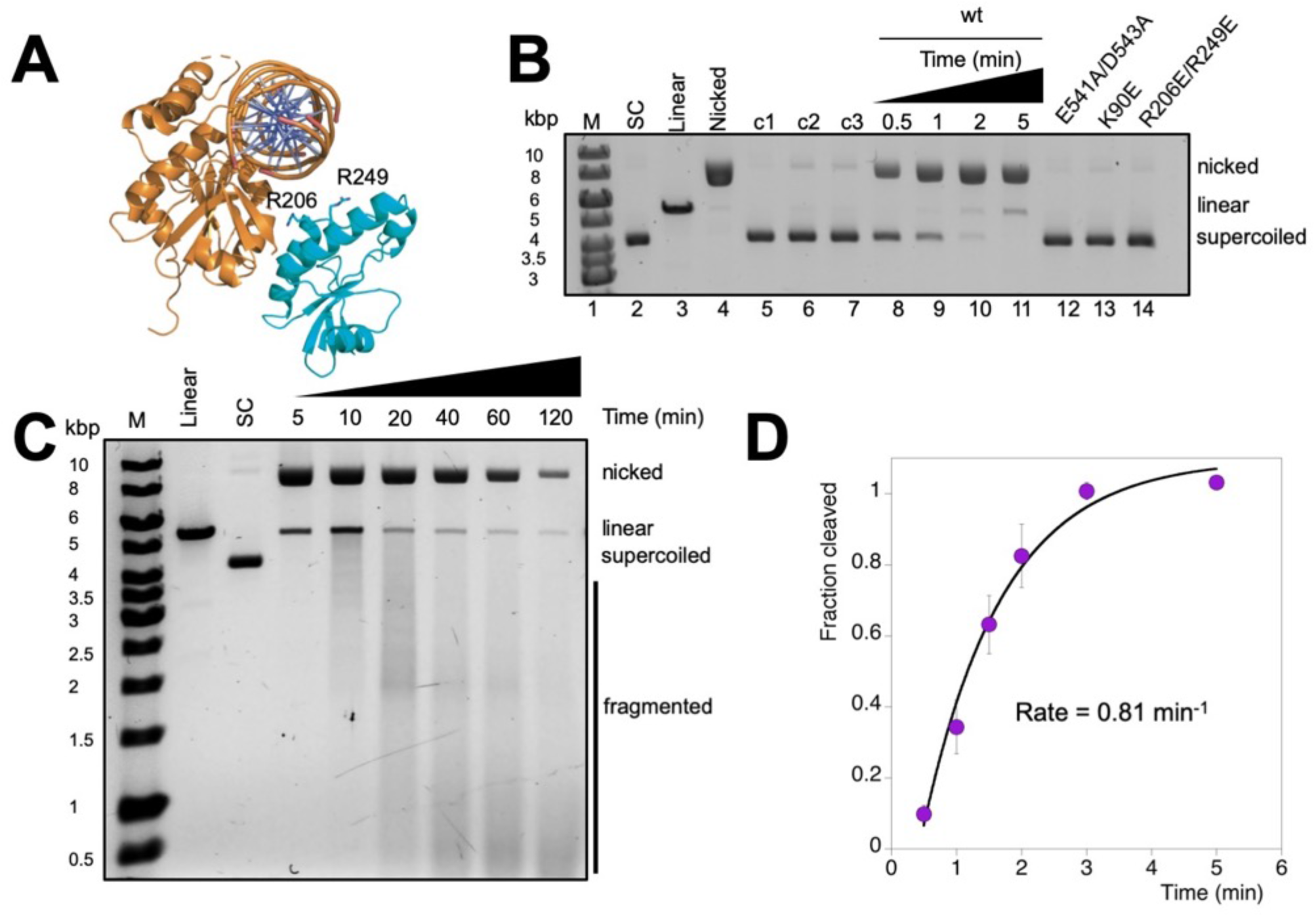
Can1 nicks supercoiled plasmid when activated by cA_4_ in the presence of MnCl_2_. **(A)** The nuclease domain of Can1 (orange) in complex with the modelled position of dsDNA (based on homologous structures, the nuclease domains of *AgeI* (PDB: 5DWB) and *NgoMIV* (PDB: 1FIU)). The nuclease-like domain of Can1 is shown in cyan and Arg206 and Arg249 are shown as sticks. **(B)** Native agarose gel analysis of plasmid (2 nM) cleavage by Can1 (200 nM), showing rapid nicking in the presence of cA_4_ (200 nM) and MnCl_2_ (5 mM) (lanes 8-11). Standards corresponding to supercoiled, linear and partially nicked plasmid are shown in lanes 2-4. Lanes 5-7 show the reactions incubated for 5 min without protein (c1), MnCl_2_ (c2) or cA_4_ (c3), respectively. Can1 variants E541A/D543A, K90E and R206E/R249E generate minimal nicked product after 5 min (lanes 12-14). M – DNA size markers. **(C)** Extended time course of plasmid degradation by Can1. Rapid nicking is followed by slower degradation to small linear products over 2 h. Standards are as described in part A. **(D)** Single-turnover kinetic analysis of supercoiled plasmid nicking by Can1, under the conditions in A, yield a rate constant of 0.81 ± 0.15 min^−1^.

Perhaps more surprisingly, DALI searches revealed that the top hit for homologues of domain 2 of Can1 is also VC1899, which overlaps with an RMSD of 2.5 Å over 86 residues. Domain 2 of Can1 also superimposes with the nuclease domain of Can1 with an RMSD of 2.6 Å over 83 residues, but it lacks the canonical nuclease active site residues of either (Figure S7). We herewith refer to domain 2 as the “nuclease-like domain”.

### Can1 is a cA_4_ activated, metal dependent DNA nuclease

The variants E541A/D543A, targeting the PD-D/ExK active site, K90E, targeting a key interaction with cA_4_ in the CARF domain, and R206E/R249E, targeting a putative DNA binding site in the nuclease-like domain, were expressed and purified as for the wild-type enzyme. Wild-type Can1 (200 nM) cleaved a supercoiled plasmid substrate (2 nM) in the presence of 200 nM cA_4_ and 5 mM MnCl_2_ (Figure 4B). The majority of the plasmid was nicked within 1 min, with a small amount linearized after 5 min incubation. Reactions in the absence of MnCl_2_ or cA_4_ showed very little activity, confirming that Can1 is a metal dependent DNA nuclease activated by cA_4_. Moreover, variants E541A/D543A (nuclease variant) and K90E (CARF domain variant) were both catalytically inactive, as expected for a cA_4_-activated PD-D/ExK family nuclease. The R206E/R249E variant, which reverses two positive charges on the surface of the nuclease-like domain, was also inactive, suggesting a role for this domain in DNA binding. These residues are in reasonably close proximity to the predicted position of the dsDNA (Figure 4A), and the nuclease-like domain may move closer still upon DNA binding.

On extended incubation over 2 h, Can1 slowly degraded the nicked plasmid to linear fragments of progressively smaller size (Figure 4C). Thus, the enzyme favours nicking of supercoiled DNA, with more extensive DNA degradation seen only under extended incubation. To measure the rates of supercoiled plasmid nicking under pseudo-single turnover conditions, 200 nM protein was incubated with 2 nM plasmid, and the fraction of supercoiled plasmid cleaved was quantified, normalised, plotted and fitted to an exponential equation as described in the Methods. Two biological replicates and six technical replicates were carried out, yielding a rate of 0.81 ± 0.15 min^−1^ (Figure 4D).

### Can1 cleaves supercoiled DNA at random sites to generate ligatable nicks

To determine whether Can1 nicks supercoiled DNA at a specific site, we compared its activity to the Nickase Nt.*Bsp*QI, which nicks DNA specifically at 5’-GCTCTTCN/ sites, and has one recognition site in the plasmid pEV5HisTEV. By using alkaline agarose gel electrophoresis to analyse the cleavage products, we ensured that DNA migrated in single stranded form. Nt.*Bsp*QI nicking generated a single product band of 5.4 kb, as expected (Figure 5A), whilst cleavage by Nt.*Bsp*QI followed by *Bam*HI (which also has a single recognition site in pEV5HisTEV) yielded 3 bands – a 5.4 kb band representing the strand that is cut only by *Bam*HI, and two smaller bands of 3.0 and 2.4 kb that result from cleavage of the Nt.*Bsp*QI nicked strand by *Bam*HI, as shown in the schematic. Can1 also generated a nicked strand of 5.4 kb, but when this product was further digested by *Bam*HI a smear of smaller DNA products were observed rather than specific fragments (Figure 5A). This demonstrates that Can1 nicks supercoiled DNA at random sites.

**Figure 5.**
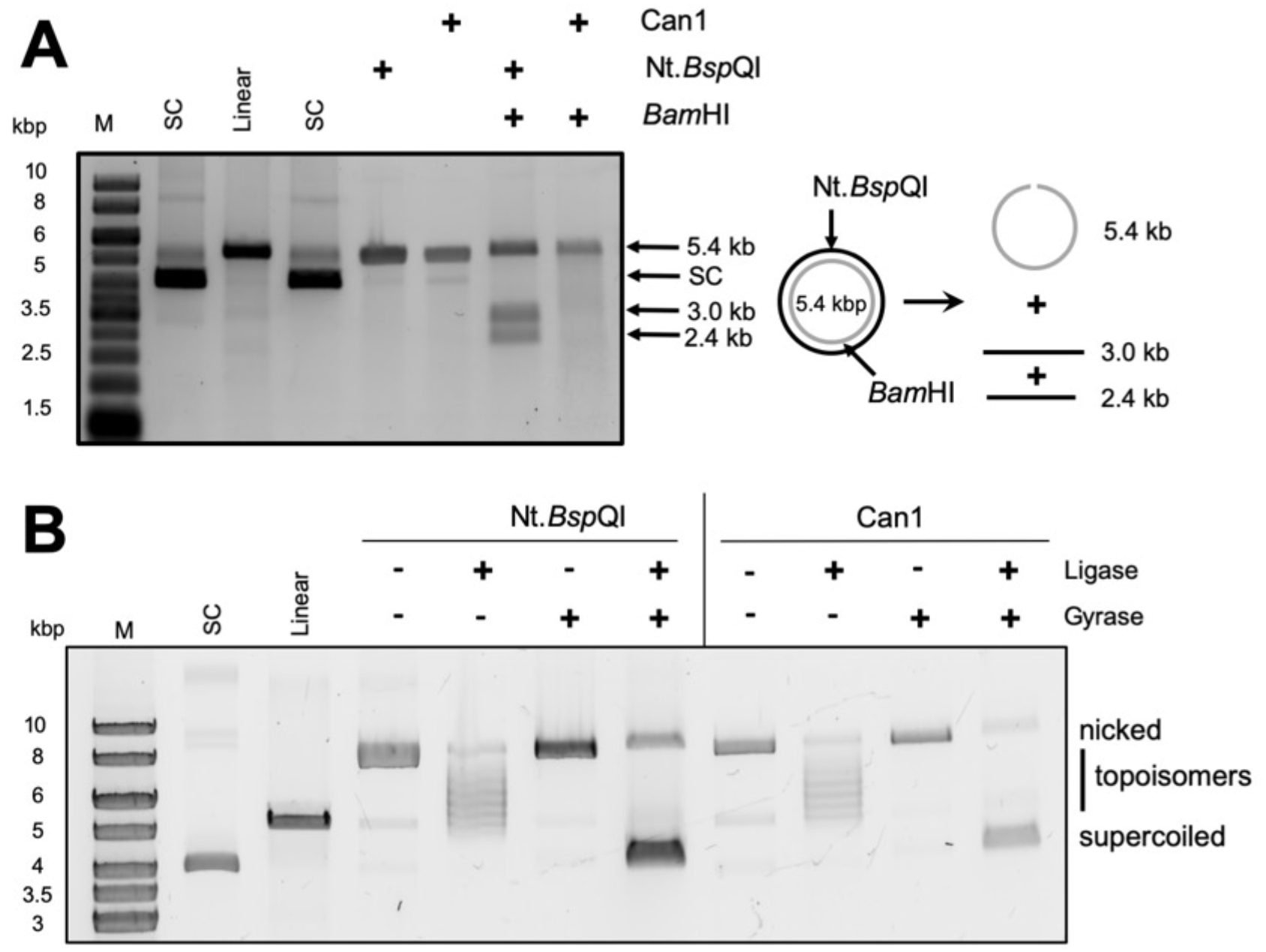
Can1 nicks plasmid DNA at random sites, leaving ligatable DNA ends. **(A)** Plasmid nicking by the Nickase Nt.*Bsp*QI and Can1 were analysed by alkaline agarose gel electrophoresis where all DNA is denatured into single-stranded products. Both generate linear 5.5 kb products. When Nt. *Bsp*QI nicking at its recognition site is followed by *Bam*HI digestion, the expected linear products of 3.0 and 2.5 kb are also observed. Can1, which nicks at random sites, generates a wide range of linear products following BamHI or Nt.*Bsp*QI digestion that appear as a smear in the gel. **(B)** Both the Nickase Nt.*Bsp*QI and Can1 nick supercoiled plasmid to yield open circle DNA that can be visualised following agarose gel electrophoresis. Ligation of these products by DNA ligase generates closed circular DNA with a range of topoisomers. Addition of DNA gyrase restores the supercoiled DNA substrate. Control lanes are as for figure 4.

Nucleases within the superfamily which contains the PD-D/ExK motif typically catalyse phosphodiester bond cleavage by a two metal ion catalytic mechanism^26^. A water molecule is deprotonated by one metal ion and used as a nucleophile to attack the scissile phosphate bond, generating 5’-phosphate and 3’-hydroxyl products^26^. To determine whether Can1 conformed to this mechanism, we incubated the nicked plasmid generated by Can1 with DNA ligase, followed by DNA gyrase, as described in the Methods. Commercial nicking endonuclease Nt.*BspQI* was used as a positive control. After incubating nicked plasmid with ligase, various relaxed topoisomers were observed (Figure 5B), suggesting that the nicked plasmid was ligated to covalently-closed circular plasmid. The products were further incubated with DNA gyrase to introduce negative supercoils (Figure 5B), confirming that ligation of the nicked sites had occurred. Thus, Can1 hydrolyses phosphodiester bonds in supercoiled DNA substrates, generating ligatable 5’-phosphate and 3’-hydroxyl products.

### Structural rearrangements upon cA_4_ binding

The binding of a ligand as large as cA_4_ will inevitably require structural rearrangements of Can1, particularly as it is completely buried in the complex. As we lacked a crystal structure of apo Can1, we investigated the solution state of apo Can1 and Can1 bound to cA_4_ using small angle X-ray scattering (SAXS). Size exclusion chromatography coupled SAXS was performed on apo Can1 to ensure optimal background subtraction and sample characterization, and a single, well defined elution peak was observed. The SAXS derived radius-of-gyration, R_g_, shows the apo state is less compact than Can1 in complex with cA_4_, as characterized by a larger R_g_ (28.44 ±0.13 Å vs 30.13 ±0.15 Å), particle volume and maximum dimension, d_max_. The dimensionless Kratky plot (Figure 6A) suggests Can1 is not globular in either the bound or apo state, which is consistent with the observations of the X-ray crystal structure of Can1 bound to cA_4_. Furthermore, analysis of the real-space pairwise distances within the particles, P(r)-distribution, shows the apo state of Can1 has a wider overall distribution (Figure 6B) with a substantially larger maximum dimension, d_max_ (121.5 Å vs 101.5 Å for the complex), suggesting there is a significant conformational difference between the bound and apo states in solution. In fact, fitting the Can1 crystal structure to the SAXS data results in a Chi^2^ > 80.

**Figure 6.**
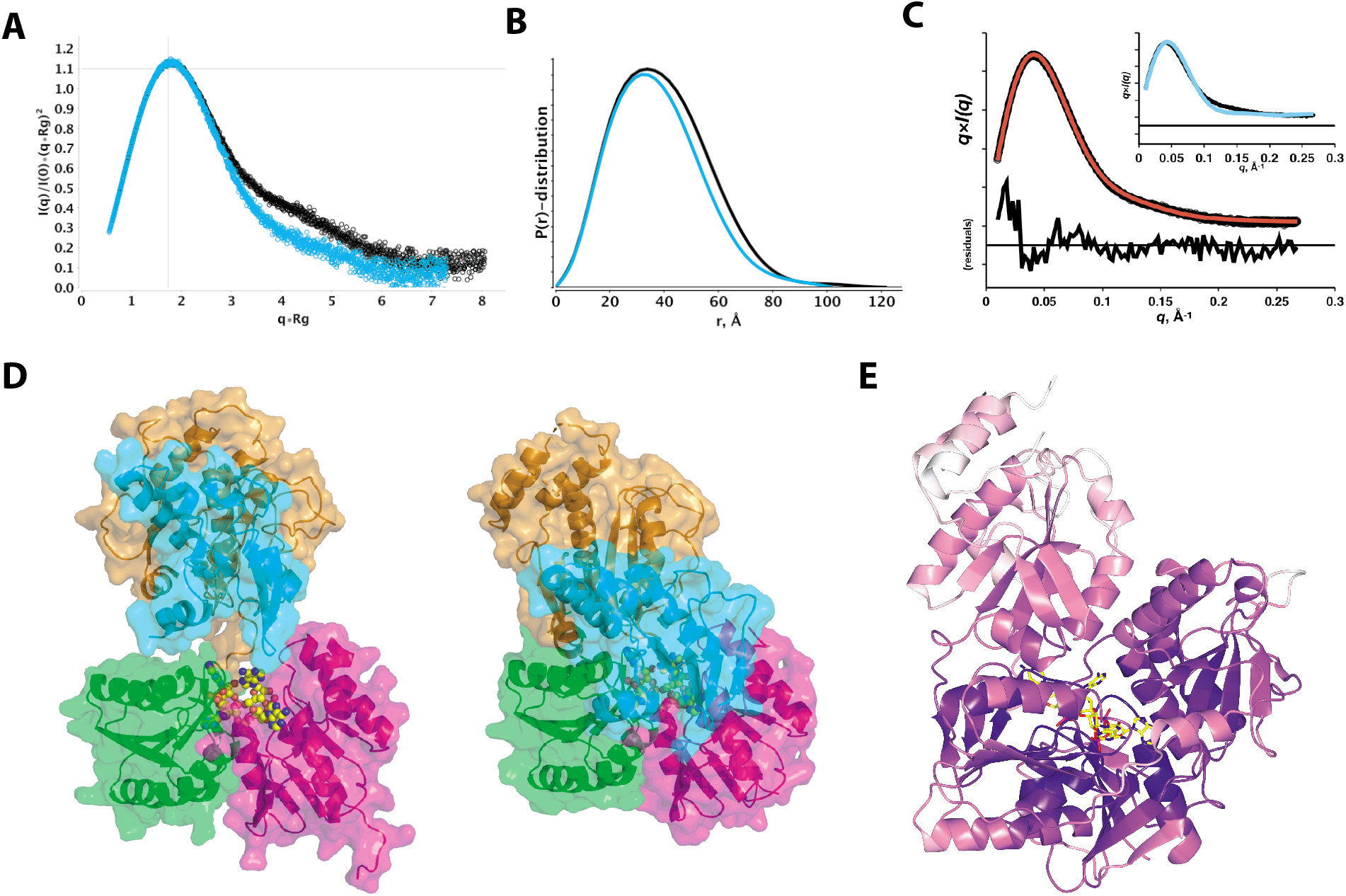
Structural rearrangements of Can1 upon cA_4_ binding in solution. **(A)** Dimensionless Kratky plot of Can1 in the presence (blue) and absence (black) of cA_4_. Data were normalized with R_g_ and I(0) of 28.88 ±0.13 Å and 1.090E-1 ±1E-4 respectively for Can1 with cA_4_ and 30.26 ±0.11 Å and 8.99E-2±6.5E-05 respectively for apo Can1. Cross-hairs mark the Guinier-Krakty point or peak position for an ideal, globular particle. **(B)** Paired-distance, P(r), -distribution function performed as an indirect Fourier transform using the Moore method. P(r)-distributions were normalized to particle’s Porod volume using 121,421 Å^3^ and 132,291 Å^3^ for the Can1 with cA_4_ (blue) and apo Can1 (black), respectively. Widening of the distribution for apo Can1 suggests larger distances are more accessible. **(C)** Fit of apo Can1 SAXS data (black circles) using atomistic model from constrained torsion angle molecular dynamics. Best single-state model (red line), *χ^2^*1.8, identifies a model that opens through translation of the nuclease domain. The best open model was achieved where the CARF dimer was maintained as a rigid body and domain 2 was allowed to vary ±5 Å from its observed position. The nuclease domain was treated as a separate, rigid body tethered by an unconstrained loop (residues 423 to 428). Residuals of the fit are presented. The inset shows the fit of the Can1 crystal structure (cyan) to apo-state SAXS data (black). **(D)** Surface representation of Can1 as predicted by SAXS in the absence of cA_4_ (left) and the structure of Can1 in complex with cA_4_ solved by X-ray crystallography (right). Coloured as first CARF domain (magenta), nuclease-like domain (cyan), second CARF domain (green), nuclease domain (orange). Loops between the domains are not shown for clarity. **(E)** The crystal structure of Can1 shown in cartoon form coloured by temperature factor from white (high temperature factor) to dark purple (low temperature factor). A molecule of cA_4_ bound to Can1 is shown in yellow stick representation.

To investigate the structural changes required to explain the SAXS dataset of the apo state, we performed a distance restrained, rigid body molecular dynamics simulation that restricted the search to the distances within the apo state P(r)-distribution. The Can1 crystal structure demonstrated clearly defined domains that were used to define rigid bodies to which centre-of-mass restraints were assigned. Given the observed conservation of the CARF dimer conformation in the cA_4_ bound Can1 and apo VC1899 structures, we treated this unit as a rigid body. We generated ~2500 conformations that were subsequently fit to the apo state SAXS data. A best single-state model was found (best χ^2^ 1.81; Figure 6C and 6D), where there is movement of the nuclease and nuclease-like domains away from the core CARF dimer. This has the effect of exposing the cA_4_ binding pocket to solvent, which is required for binding. Analysis of the temperature factor of residues in Can1 (Figure 6E) shows the nuclease domain residues have higher average temperature factors than the other domains in Can1, which supports the hypothesis that the nuclease domain is more mobile. We envisage Can1 in solution is sampling between the open and closed states, exposing the cA_4_ binding site in the CARF dimer. Upon cA_4_ binding, the closed conformation of the protein is stabilized, as observed in the crystal structure. We do not rule out the possibility that a more complex population of open and closed structures exists in the absence of cA_4_.

## Discussion

### Evolution of Can1 – gene duplication and fusion

The structure of the Can1 enzyme clearly demonstrates that the two halves of the protein, comprising a CARF domain N-terminal to a nuclease fold domain, are related to one another. Coupled to the observation that each half is clearly related to the DUF1887 family, as exemplified by the VC1889 protein, we can postulate a scenario whereby gene duplication, fusion and subsequent divergent evolution of an ancestral DUF1887 family member gave rise to the Can1 family. Can1 appears to be limited to members of the genus *Thermus*, but in contrast DUF1887 is much more widely distributed, with representatives found throughout the bacterial phyla and is particularly common in the proteobacteria. Although no member of the DUF1887 family has yet been characterised biochemically, given the relationship with Can1 and the conservation of a nuclease active site we suggest the provisional name “Can2” (CRISPR ancillary nuclease 2) for this family.

### Recognition of cA_4_

As this is the first structure available for a CARF domain protein bound to its cognate cyclic oligoadenylate ligand, it is worth drawing some general conclusions about the mechanism of recognition. Can1 is unusual in being a monomeric protein with two non-identical CARF domains, and therefore possesses less intrinsic symmetry than most family members. cA_4_ is bound in a markedly asymmetrical conformation, which is in part exaggerated by the position of one adenine base in an axial position, driven by a stacking interaction with a tryptophan residue. Many different residues, the vast majority from the two CARF domains, contribute to the ligand binding pocket, which appears to be largely pre-formed in the apo protein. A significant number of these interactions are with protein main chain atoms, and although some of these are structurally conserved between the two ‘halves’ of Can1, the difference in side chain residue meant they would not have been predicted by sequence alone. Lysine 90, which occupies a central position in CARF domain 1 at the N-terminus of an a-helix, forms a salt bridge with a bridging phosphate of cA_4_ and basic residues at an equivalent position have been shown essential for cA_4_ recognition in other CARF family proteins^10,12^.

### Activation of Can1 by cA_4_ binding

The cA_4_ ligand is completely buried in the Can1 complex, demonstrating that significant protein conformational changes are required to facilitate binding. The close structural agreement observed for the two CARF domains of Can1 compared to the VC1899 apo protein suggests that the CARF dimer conformation does not change radically upon cA_4_ binding. In contrast, the temperature factors of the residues in the crystal structure, coupled with SAXS analysis, suggest that the nuclease domain is mobile, which would help facilitate cA_4_ binding to apo Can1. Modelling of dsDNA binding by the nuclease domain, coupled with the observation that conserved basic residues in the nuclease-like domain are essential for activity (presumably due to a role in DNA binding rather than directly in catalysis), suggests that cA_4_ binding allows the nuclease and nuclease-like domains to assemble into a functional dsDNA binding site, activating the enzyme. Modelling of dsDNA from homologous structures suggests a further modest closure of the cleft between the two domains may occur upon DNA binding, although this will depend on the precise position of the dsDNA substrate bound to Can1.

### Function of Can1 in vivo

An emerging theme of CRISPR-based defence systems is the role of collateral (non-specific) nucleic acid degradation in immunity (reviewed in^31^). In addition to the cOA activated, non-specific ribonucleases described above for type III systems, type VI (Cas13) effectors provide effective immunity against mobile genetic elements by degrading RNA non-specifically^32,33^. The mechanism for this immunity was recently shown to result in degradation of host, rather than virus, RNA, driving infected cells into dormancy and thus preventing the spread of infection^34^. DNA is also a target for collateral degradation. Type III effectors hydrolyse ssDNA on binding target RNA – an activity that appears largely non-specific *in vitro*^35–37^, but which may favour transcribing phage DNA *in vivo*^38^. Likewise, type V effectors such as Cas12a possess a non-specific ssDNA degradation activity that is licensed by crRNA-dependent DNA binding^39,40^.

Here, we have described another option available to CRISPR defence systems: cyclic oligoadenylate-dependent supercoiled DNA cleavage by the Can1 enzyme. Once activated by cA_4_ generated in response to detection of foreign RNA, Can1 cleaves supercoiled DNA to generate nicked products. *In vivo*, such an activity is likely to slow down the replication kinetics of rapidly-replicating phage, where nicks can result in the collapse of DNA replication forks to generate doublestrand breaks. In contrast, such nicks are not too great a burden for the slowly-replicating host chromosome as they are easily repaired by DNA ligase. This nickase activity will operate in parallel with non-specific RNA degradation by the *T. thermophilus* Csm6 enzymes (Figure 7), providing layered defence against mobile genetic elements detected by the CRISPR system. Given the structural conservation of CARF and nuclease domains in the Can2 (DUF1887) family, we predict that this family, which is widespread in bacteria, functions in a similar way to Can1 by cleaving supercoiled DNA. However, there is a key difference – the Can2 homodimer has two nuclease active sites, which may have functional consequences. In conclusion, we have revealed key new molecular details of the cOA-mediated anti-viral defence mediated by type III CRISPR systems – one of the most common types of CRISPR effectors found in nature.

**Figure 7.**
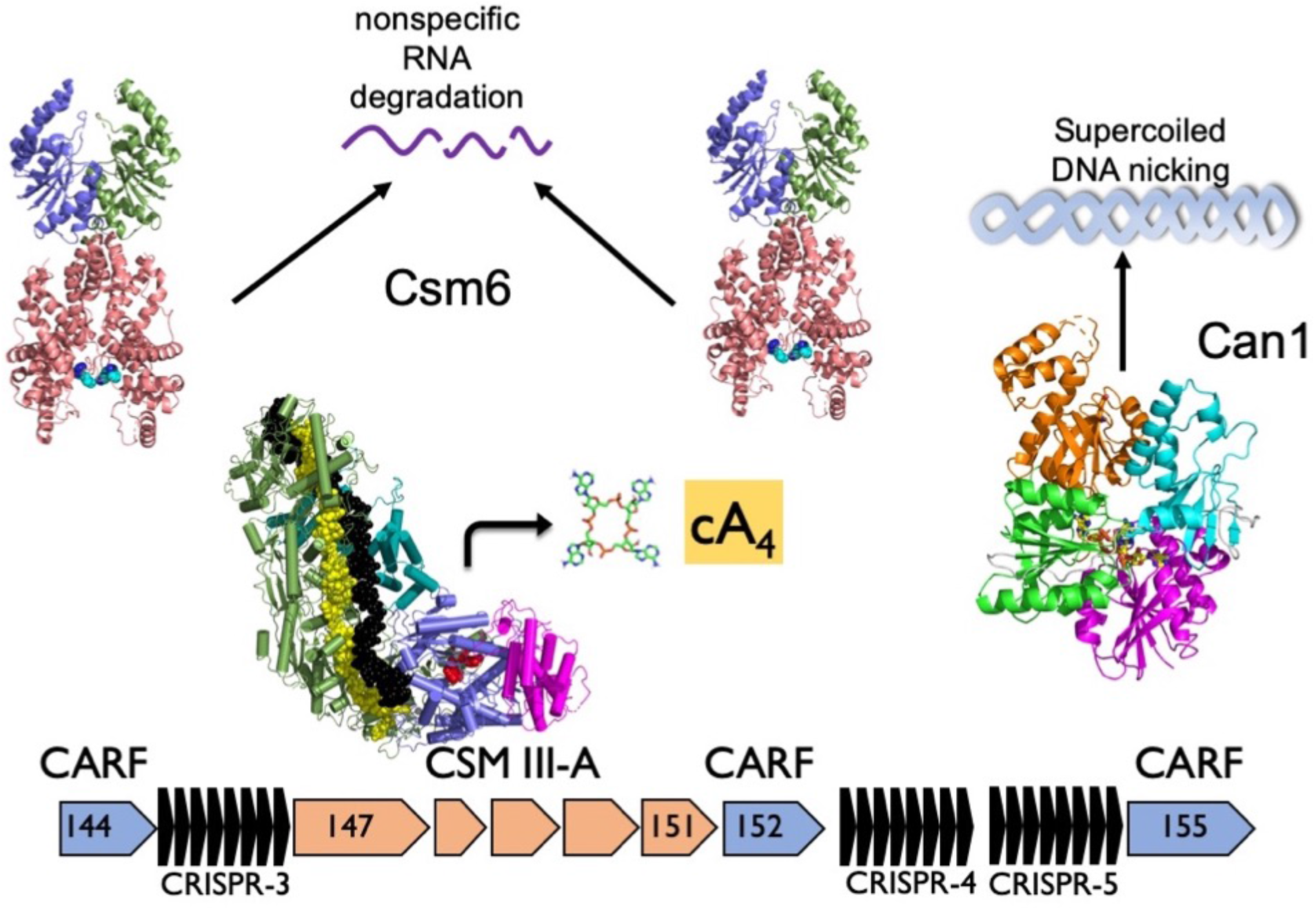
Cartoon of the type III-A CRISPR defence system of *T. thermophilus.* The type III-A effector binds phage RNA, activating the cyclase domain which synthesises cA_4_ from ATP. cA_4_ binds to and activates the two Csm6 family RNases (encoded by *tthb144* and *tthb152*) and the DNA nickase Can1 (encoded by *tthb155*), providing a multi-layered defence against mobile genetic elements.

## Materials and Methods

### Expression and purification of Can1 and variants

*can1* was purchased as a synthetic gene from Integrated DNA Technologies, Coralville, IA, United States (IDT) and cloned into the pEV5HisTEV vector^25^. The cloning strategy resulted in the addition of two residues (Leu and Glu) to the C-terminus of the protein. The full sequence of the synthetic gene is included in the supplementary materials. The variants E541A/D543A, K90E, R206E and R249E were generated using the QuikChange Site-Directed Mutagenesis kit as per manufacturer’s instructions (Agilent Technologies). The pEV5HisTEV-*can1* wild-type and variant constructs were transformed into C43 (DE3) *E. coli* cells. Protein expression was induced with 0.4 mM isopropyl-β-D-1-thiogalactoside (IPTG) at an OD_600_ of ~0.6 and grown overnight at 25 °C. Cells were harvested and resuspended in lysis buffer containing 50 mM Tris-HCl pH 7.5, 0.5 M NaCl, 10 mM imidazole and 10% glycerol, and lysed by sonicating six times for 2 min on ice with 2 min rest intervals. Can1 was purified with a 5 ml HisTrapFF column (GE Healthcare), washing with 5 column volumes (CV) of buffer containing 50 mM Tris-HCl pH 7.5, 0.5 M NaCl, 30 mM imidazole and 10% glycerol, and eluting with a linear gradient of buffer containing 50 mM Tris-HCl pH 7.5, 0.5 M NaCl, 0.5 M imidazole and 10% glycerol across 15 CV. Protein containing fractions were concentrated and the hexa-histidine affinity tag was removed by incubating protein with Tobacco Etch Virus (TEV) protease (10:1) overnight at room temperature.

Cleaved Can1 was isolated from the TEV protease by repeating the immobilised metal affinity chromatography step and collecting the unbound fraction. Size exclusion chromatography was used to further purify Can1, eluting protein isocratically with buffer containing 20 mM Tris-HCl pH 7.5, 150 mM NaCl. The protein was concentrated using a centrifugal concentrator, aliquoted and frozen at – 80°C. For seleno-methionine labelling, the pEV5HisTEV-Can1 construct was transformed into B834 (DE3) *E. coli* and cultures grown in M9 minimal medium supplemented with Selenomethionine Nutrient Mix (Molecular Dimensions, Newmarket, Suffolk, UK) and 50 mg L^−1^ (L)-selenomethionine (Acros Organics). Protein was purified as described above.

### Can1 nuclease assay

200 nM protein monomer was incubated with 2 nM plasmid pEV5HisTEV for the time indicated at 60 °C in 20 μl final reaction volume in 20 mM MES pH 6.5, 100 mM NaCl and 1 mM EDTA supplemented with 5 mM MnCl_2_ and 200 nM cA_4_. Reactions were stopped by addition of 10 mM EDTA. Control reactions were carried out by incubating plasmid without protein, MnCl_2_ or cA_4_ under the same conditions. For kinetic analysis, triplicate experiments were carried by incubating wild type Can1 and its variants in 140 μl final reaction volume under the conditions described above. 20 μl was removed and quenched by addition of 2 μl 100 mM EDTA at desired time points. After adding 4 μl 6x DNA loading dye (New England BioLabs, Ipswich, MA, United States), 10 μl sample was analysed by 0.7% native agarose gel electrophoresis. Gels were scanned by Typhoon FLA 7000 imager (GE Healthcare) at a wavelength of 532 nm, quantified using the Bio-Formats plugin^41^ of ImageJ as distributed in the Fiji package^42^ and plotted against the time using Kaleidagraph (Synergy Software, Reading, PA, United States). The data were fitted to a single exponential curve as previously described^1^.

### Nicking endonuclease digestion

Nicking endonuclease Nt.*BspQI* (New England BioLabs, Ipswich, MA, United States) hydrolyses the phosphodiester bond to 3’-hydroxyl and 5’-phosphate on one strand of dsDNA. Therefore, open circle conformation of pEV5HisTEV containing the unique GCTCTTCN^Λ^ site was generated by Nt.*BspQI*. All incubations with commercial enzymes followed manufacturer’s instructions.

### Alkaline agarose gel electrophoresis

DNA products were separated on 0.7 % alkaline agarose gels (30 mM NaCl, 2 mM EDTA and 0.7% agarose). The gels were immersed for 2 h in alkaline electrophoresis buffer (30 mM NaOH, 2 mM EDTA) before running. Samples and ladder were denatured with 6x alkaline electrophoresis loading dye (180 mM NaOH, 6 mM EDTA, 18% Ficoll 400 and 0.05% bromophenol blue) and heated at 95 °C for 5 min, then chilled on ice. Gels were run at 2 V/cm for 10 h in alkaline electrophoresis buffer, then soaked in renaturing buffer (500 mM Tris-HCl pH 8.0) for 30 min and stained with SYBR Gold (Thermo Scientific, Waltham, MA, United States) for 30 min. Gels were destained with H_2_O and scanned by Typhoon FLA 7000 imager at a wavelength of 532 nm.

### Plasmid ligation and supercoiling

200 nM protein was incubated with 2 nM plasmid pEV5HisTEV in 20 μl final reaction volume as described above. Reactions were quenched and deproteinized by PCR Clean-Up System (Promega, Madison, Wisconsin, United States). 10 μl eluted product was incubated with T4 DNA Ligase (New England BioLabs, Ipswich, MA, United States) in ligase buffer (50 mM Tris-HCl pH 7.5, 10 mM MgCh, 1 mM ATP and 10 mM DTT) in 20 μl final reaction volume following manufacturer’s instructions. After inactivating at 65 °C for 10 min, ligated products were incubated with *E. coli* DNA gyrase (Inspiralis, Norwich, UK) in in gyrase buffer (35 mM Tris-HCl pH 7.5, 24 mM KCl, 4 mM MgCh, 2 mM DTT, 1.8 mM spermidine, 1 mM ATP, 6.5% glycerol and 0.1 mg/ml albumin) in 40 μl final reaction volume as per manufacturer’s instructions. Reactions were deproteinized by phenol chloroform extraction before analysis by agarose electrophoresis.

### Crystallisation of Can1

Selenomethionine labeled Can1 at 10.4 mg/ml was mixed in a 1:2 molar ratio with cA_4_ and incubated at room temperature for 30 minutes, before centrifugation at 13,000 rpm prior to crystallization. Sitting drop vapour diffusion experiments were set up at the nanoliter scale using both commercially available and in-house crystallization screens, and subsequently incubated at 293 K. Crystals appeared in many conditions, but those used for data collection grew from a reservoir solution of 20% PEG 3350, 0.2 M sodium citrate, and 0.1 M bis-Tris propane, pH 6.5. Crystals were harvested and transferred to a drop of reservoir solution with the addition of 20% glycerol before cryo-cooling. Attempts to crystallize Can1 in the absence of cA_4_ did not yield crystals.

### X-ray data collection, data processing, structure solution, and refinement

X-ray data were collected at a wavelength of 0.9159 Å, on beamline I04-1 at Diamond Light Source, at 100 K to 1.74 Å resolution. Data were automatically processed using Xia2^43^ using XDS and XSCALE^44^. A strong anomalous signal was detected to 2.36 Å resolution. The data were phased using AutoSol in Phenix and the initial model was built in AutoBuild^45^. Model refinement was achieved by iterative cycles of REFMAC5^46^ in the CCP4 suite^47^ and manual manipulation in COOT^48^. Electron density for cA_4_ was clearly visible in the maximum likelihood/σ_A_ weighted *F*_obs_ – *F*_calc_ electron density map at 3σ. The coordinates for cA_4_ were generated in ChemDraw (Perkin Elmer) and the library was generated using acedrg^27^, before fitting of the molecule in COOT. Model quality was monitored throughout using Molprobity^49^. Data and refinement statistics are shown in Table 1. The coordinates and data have been deposited in the Protein Data Bank with deposition code ****.

**Table 1:** Data collection and refinement statistics for Can1 in complex with cA_4_

|  |  |
| --- | --- |
| **Data Collection** |  |
| Space group | P 2<sub>1</sub> 2<sub>1</sub> 2<sub>1</sub> |
| Cell dimensions |  |
| a, b, c (Å) | 84.4, 84.5, 123.1 |
| α, β, γ (°) | 90, 90, 90 |
| Resolution (Å) | 59.7 – 1.83 (1.86 – 1.83)* |
| R<sub>merge</sub> | 0.07 (1.43)* |
| I/σ(I) | 18.0 (2.0)* |
| Completeness (%) | 100 (100)* |
| Anomalous completeness | 100 (99.9)* |
| Multiplicity | 13.3 (13.4)* |
| Anomalous multiplicity | 6.9 (6.9)* |
| CC<sub>1/2</sub> | 0.999 (0.755)* |
| V<sub>m</sub> (Å<sup>3</sup>/Da) | 3.05 |
| Solvent content (%) | 59.7 |
| Mid-Slope of anomalous normal probability | 1.20 |
| **Refinement** |  |
| Unique reflections | 78941 |
| R<sub>work</sub> / R<sub>free</sub> | 17.7 / 21.1 |
| RMSD bonds (Å) | 0.010 |
| RMSD angles (°) | 1.56 |
| Number of atoms (non hydrogen): |  |
| Protein | 4884 |
| Water | 415 |
| cA<sub>4</sub> | 88 |
| B factors (Å<sup>2</sup>): |  |
| Protein | 42.5 |
| Water | 46.9 |
| cA<sub>4</sub> | 25.4 |
| Ramachandran plot: |  |
| Favoured / outlier (%) | 98.5 / 0.0 |
| Molprobity score / centile | 1.05 / 100 |

\* Values in parentheses refer to the high resolution shell
RMSD, root mean square deviation

### Small angle X-ray scattering

The apo-state of Can1 was measured using size exclusion chromatography coupled to small angle X-ray (SEC-SAXS) on beamline B21 at the Diamond Light Source. A 4.8 ml Shodex KW-402.5 column was pre-equilibrated with 3 column volumes of buffer containing 20 mM Tris pH 7.5 and 150 mM NaCl. SEC-SAXS experiments were measured at a flow rate of 0.160 ml per minute with 2 second exposures per frame in a 1.5 mm diameter, 10 micron thick quartz capillary flow cell and a 800 micron squared X-ray beam focused on an Eiger 4M detector 4 meters from the sample. Column performance and instrumentation calibration was checked by measuring a 45 μl sample of BSA injected at 10 mg/ml. 45 μl of the apo-state of Can1 was injected (10.4 mg/ml, 140 μM) and frames were collected for a total of 30 minutes (Supplementary information). The flat radius-of-gyration, R_g_, and nearly ideal overlay of the frames constituting the merged frames suggests the measured frames are free of interparticle interference (Supplementary information).

To measure Can1 in the presence of cA_4_, a SEC purification of Can1 was performed at the beamline by manually collecting the peak fraction (~100 μl) during an additional SEC run of the same 10.4 mg/ml protein stock. The peak fraction represented a dilution of ~3.2x for a final concentration of 44 μM of purified protein. A corresponding buffer blank (~1 ml) was collected before sample injection. Samples were mixed with ligand to a final concentration of 0.25 mM and 3 dilutions of the sample were made first by making a two-thirds dilution of the master stock followed by two consecutive nine-tenths dilutions. Dilutions were performed with the buffer blank supplemented with ligand to maintain a ratio of 3-to-1 ligand-to-protein ratio. Samples were processed in batching mode at 25 μl per sample using the Arinax sample handling robot.

Raw SAXS images were processed with the DAWN^50^ processing pipeline at the beamline to produce normalized, integrated 1-D un-subtracted SAXS curves. SEC-SAXS analysis and buffer subtractions were performed with the program ScÅtter (www.bioisis.net). Modeling of the apo-state was performed using the program CNS version 1.3 (cns-online.org/v1.3)^51^. First, missing loops and tails from the X-ray crystal structure were added by building an extended chain model of Can1 with generate_extended.inp. Protein only distance restraints were extracted from the ligand bound crystal structure of Can1 and used as pseudo-NOE constraints to fold the extended chain into the known parts of the crystal structure (model_anneal.inp). The resulting structure matched the crystal structure with missing loops and tails added back but in unconstrained positions. The folded structure (ground-state) was used as a starting conformation for limited rounds of simulated annealing or constant temperature simulations (anneal.inp) to sample various conformations as domain-to-domain distance restraints were varied. Specifically, the two CARF domains were kept as a rigid unit, whereas either the nuclease or nuclease-like domains were allowed to vary from their ground-state positions as rigid bodies. An initial simulation allowed the non-CARF domains to move as unconstrained rigid bodies only tethered by the linking peptide regions. Additional simulations were performed that constrained the non-CARF domains within 5 and 10 Å of their observed crystallographic positions. These were loose restraints added between the centre-of-masses for each domain based on the P(r)-distribution of the apo-state SAXS data. Specifically, no distances were allowed to exceed *d_max_* and distances were given upper and lower limits determined as *π/q_max_*. A total of 2430 conformations were generated and used as an ensemble fit with the online web-app multi-FOXS^52^ which identified the best two-state model for the apo-state. Single model fits were performed with FOXS (http://modbase.compbio.ucsf.edu/foxs)^53^.

## ACKNOWLEDGEMENTS

Thanks to the University of St Andrews Mass Spectrometry facility, and to staff at Diamond Light Source (beamlines I04-1 and B21).

## Supplementary Figures

**Figure S1.**
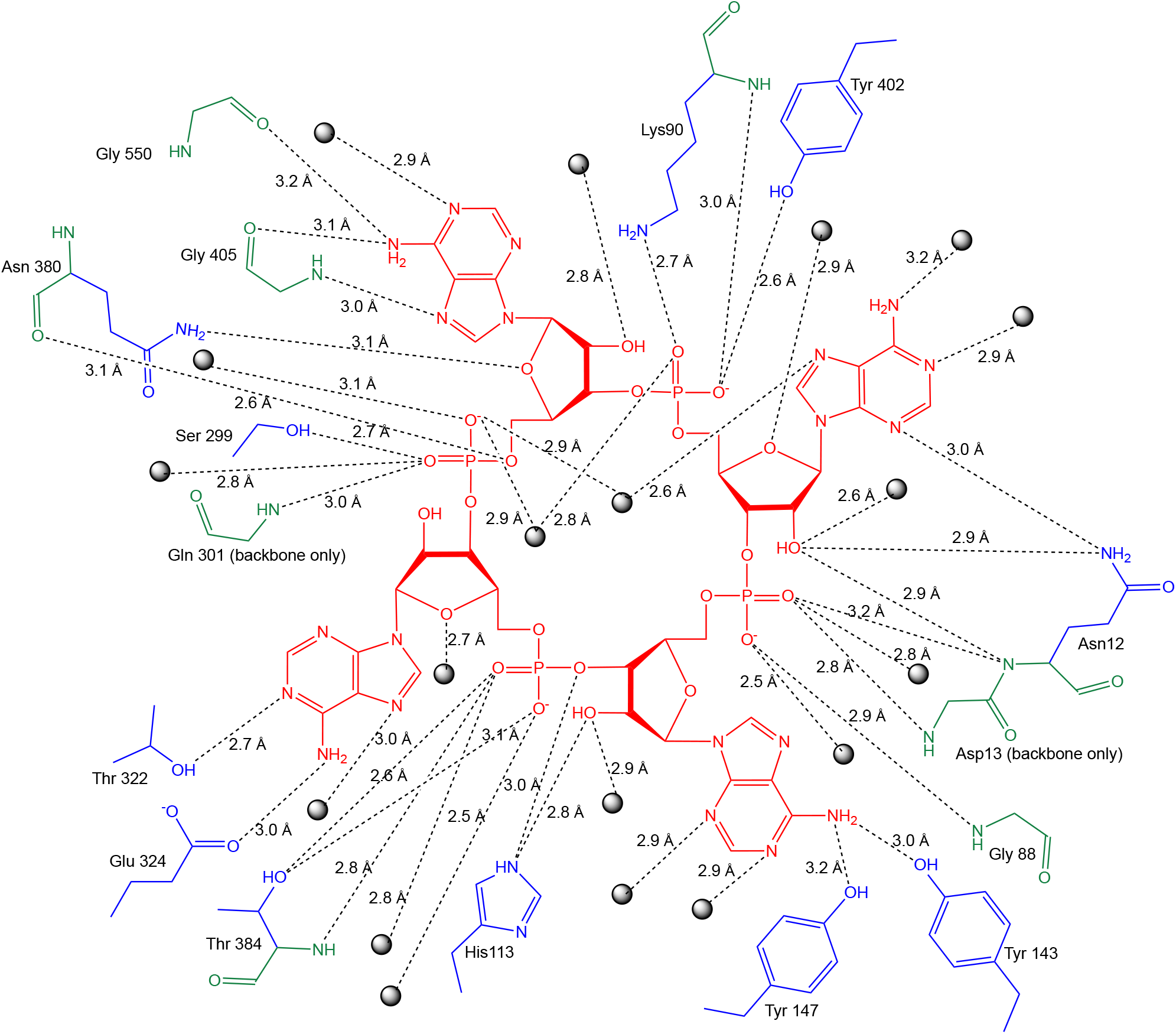
Schematic showing interactions between Can1 and cA_4_. cA_4_ is shown in red, interactions with main chain atoms in green, interactions with side chain atoms in blue, and water molecules are represented as spheres. Hydrogen bonds are represented as dotted lines, annotated with the distance.

**Figure S2.**
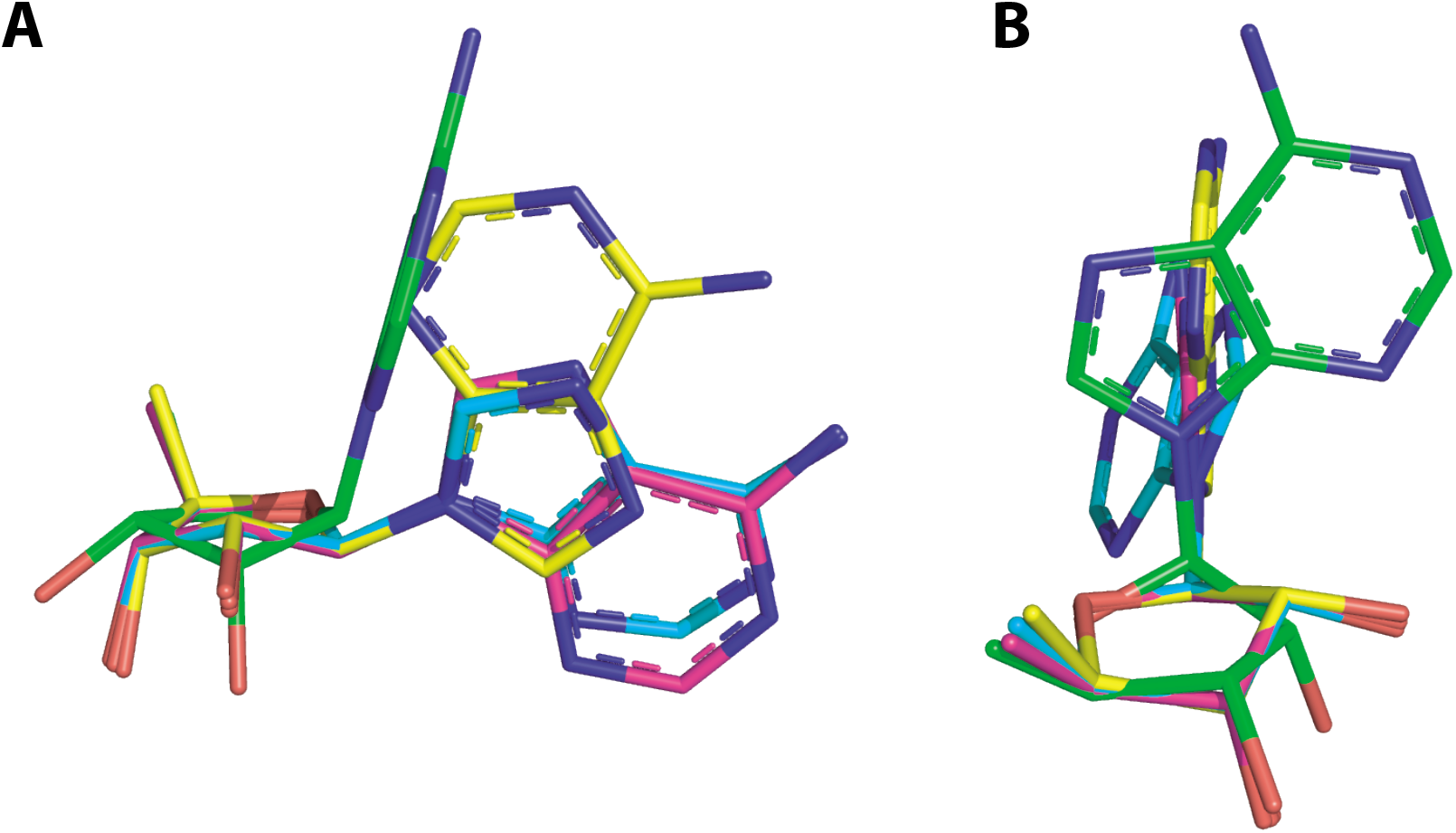
Superposition of the four adenosine units in cA_4_. The superposition is shown from two different views (**A** and **B**), with each adenosine unit indicated by a differ colour. There is a marked difference in the conformation of the ribose ring and position of the adenine base for one of the units (shown in green) which is brought about by a π-π stacking interaction with Trp42.

**Figure S3.**
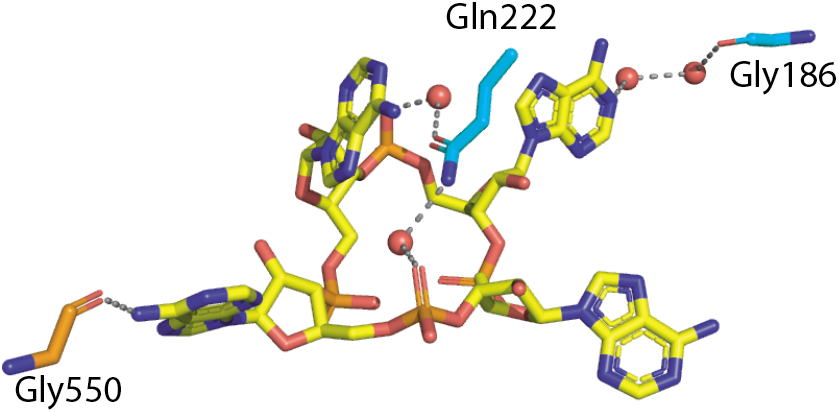
Interactions of non-CARF domain residues with cA_4_. cA_4_ and residues are shown in stick representation, with cA_4_ in yellow and amino acid residues coloured by the domain from which they originate (domain 2: cyan; nuclease domain: orange). The dotted lines represent hydrogen bond interactions, and red spheres represent water molecules.

**Figure S4.**
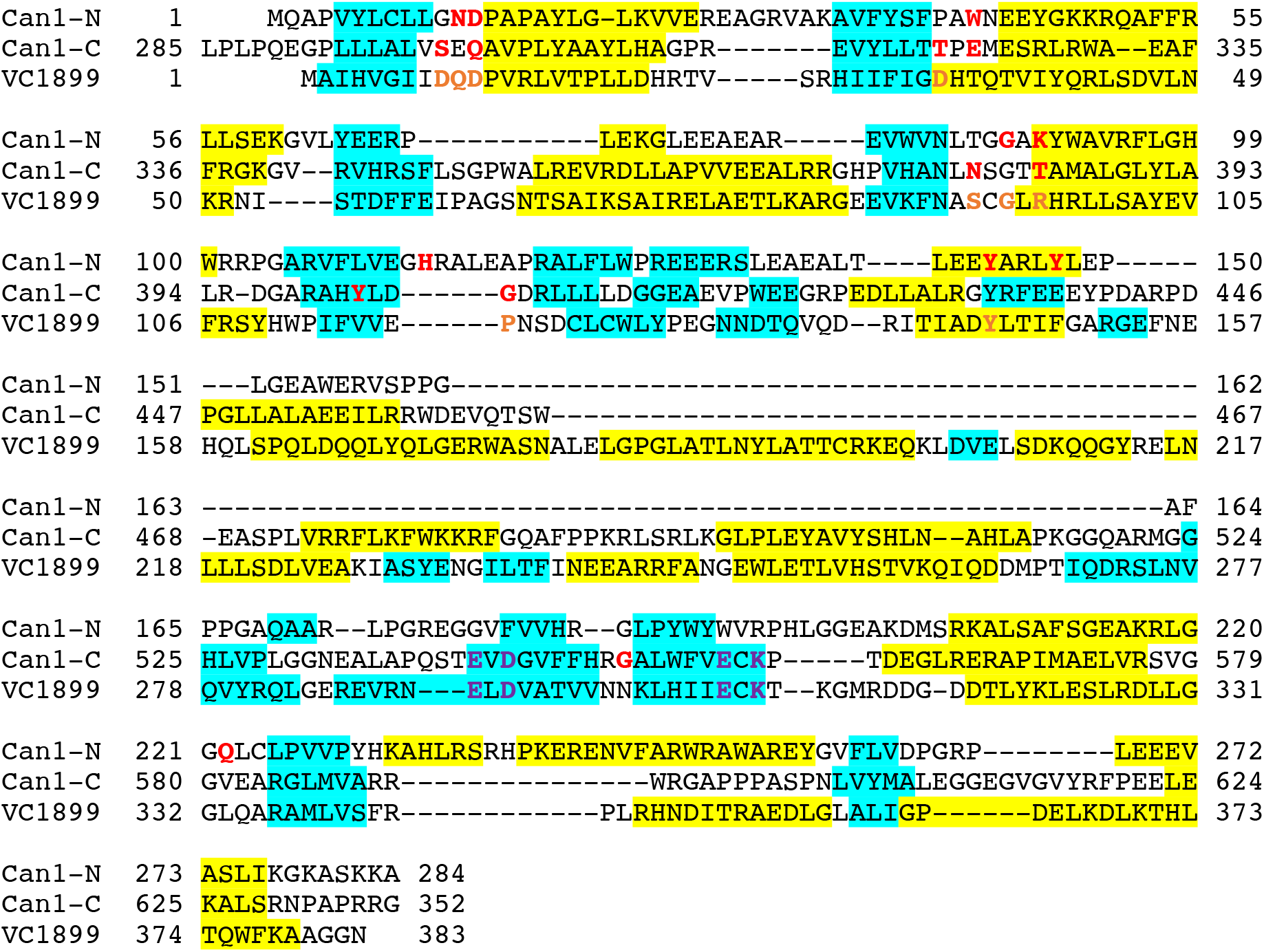
Sequence alignment based on structural homology of the first CARF domain and the nuclease-like domain of Can1 (Can1-N), the second CARF domain and nuclease domain (Can1-C) and VC1899. α-helices are highlighted in yellow and β-strands in cyan. Residues in Can1 that interact with cA_4_ are coloured red, and predicted cA_4_ interacting residues in VC1899 are coloured orange. Nuclease active site residues are coloured purple.

**Figure S5.**
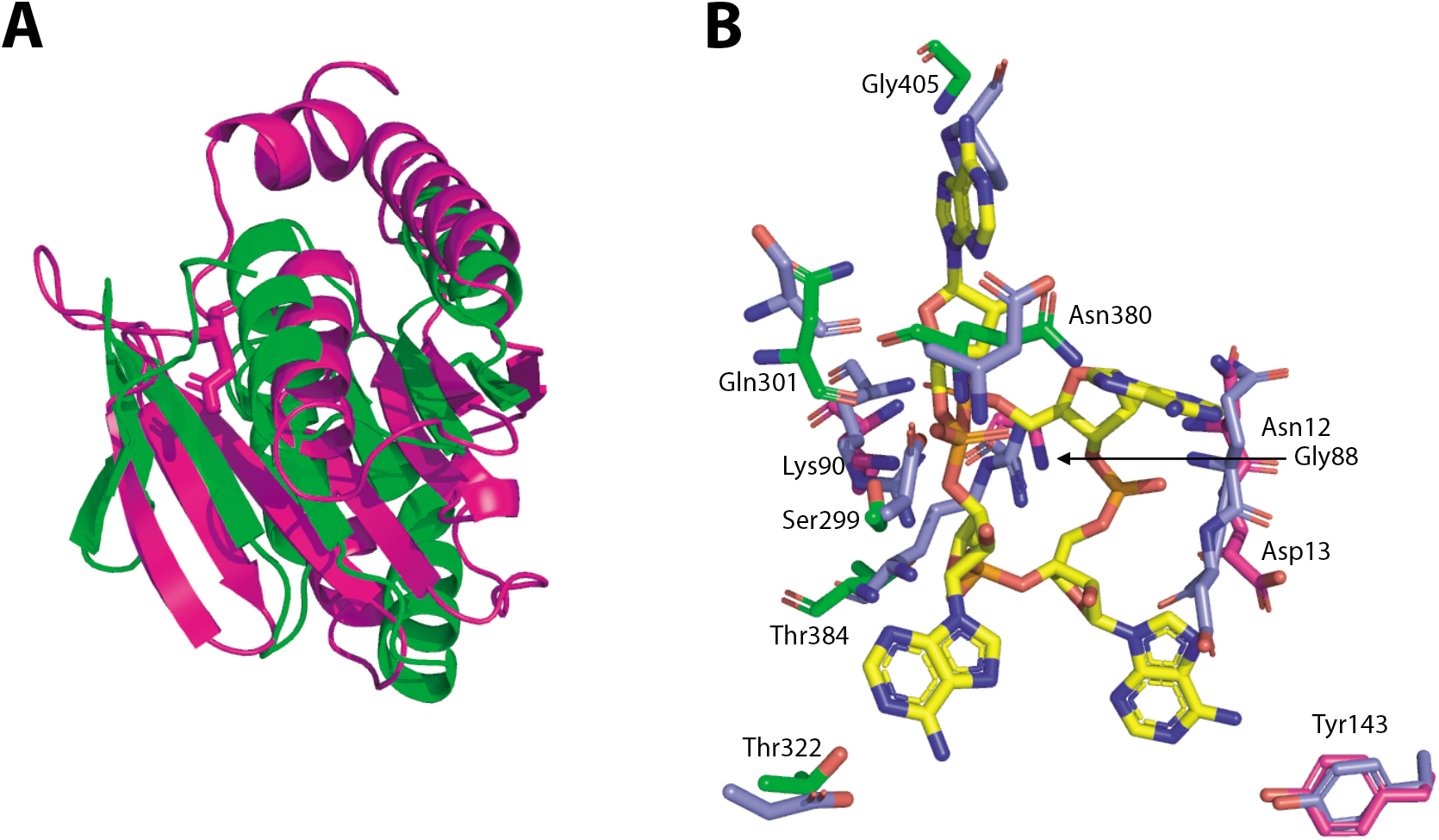
**(A)** Superimposition of the two CARF domains of Can1. The first CARF domain is shown in magenta and the second CARF domain in green. **(B)** The binding site of cA_4_ in complex with Can1, overlapped with VC1899. cA_4_ and binding site residues are shown in stick representation, with cA_4_ in yellow and amino acid residues coloured by the domain from which they originate (CARF domain 1: magenta, CARF domain 2: green). VC1899 (mauve) is superimposed onto the CARF domain of each ‘half’ of Can1, and those residues that are structurally conserved are shown. Numbering in the figure refers to residues from Can1. The equivalent residues in VC1899 are: Arg96 to Lys90 (CARF1) and Thr384 (CARF2), Gln10 to Asn12, Asp11 to Asp13 (CARF1) and Gln301 (CARF2), Gly94 to Gly88, Pro118 to Gly405, and Ser92 to Asn380 (all main chain interactions), and Arg96 to Lys90 (CARF1) and Thr384 (CARF2), Gln10 to Asn12, Tyr145 to Tyr143, Asp34 to Thr322, Asp9 to Ser 299, and Ser 92 to Asn380 (all side chain interactions).

**Figure S6.**
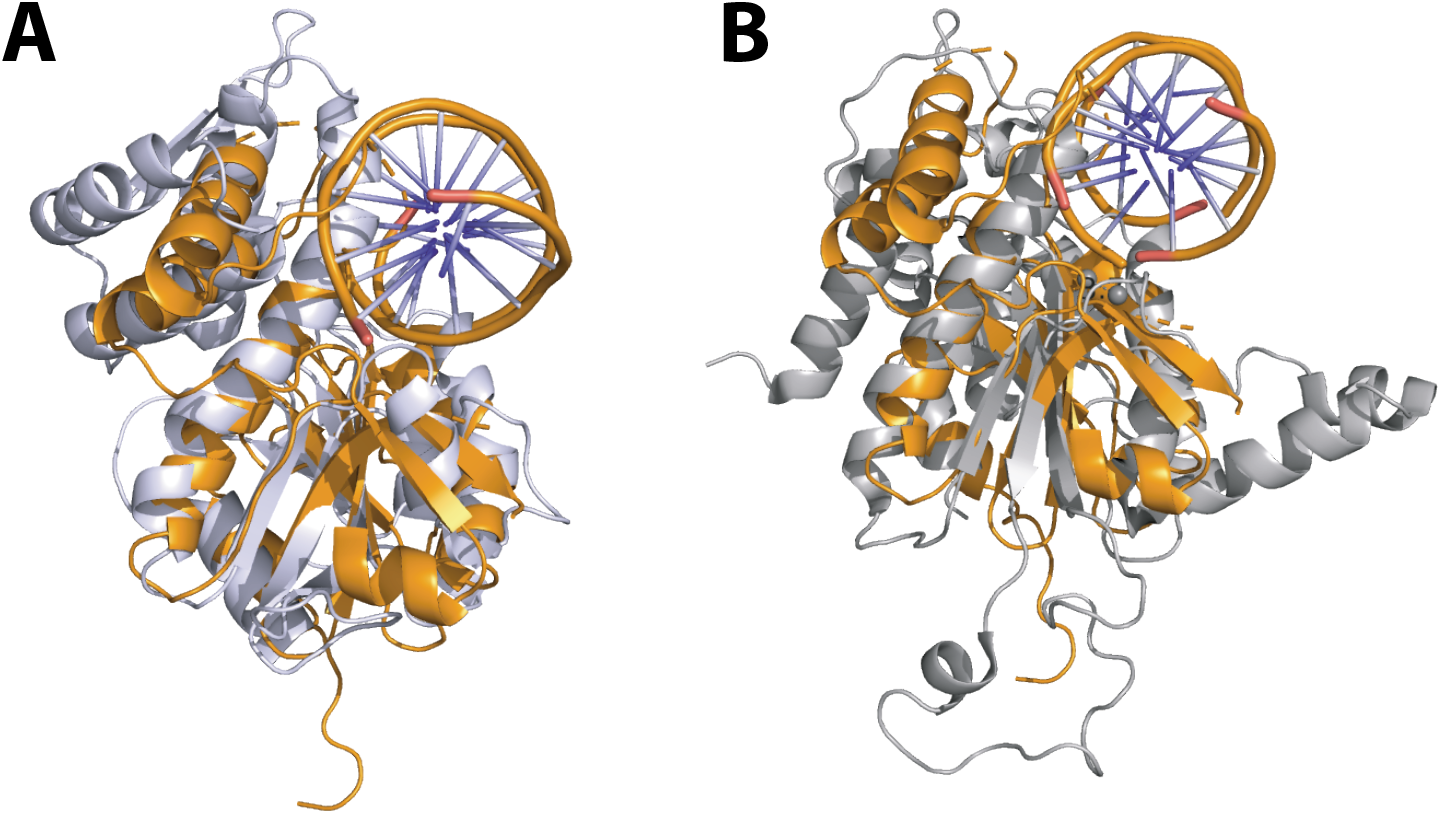
DNA binding models for Can1. The nuclease domain of Can1 (orange) overlapped with **(A)** the nuclease domain of *AgeI* (PDB: 5DWB) (light mauve) in complex with dsDNA shown in cartoon form and **(B)** the nuclease domain of NgoMIV (PDB: 1FIU (light grey) in complex with dsDNA.

**Figure S7.**
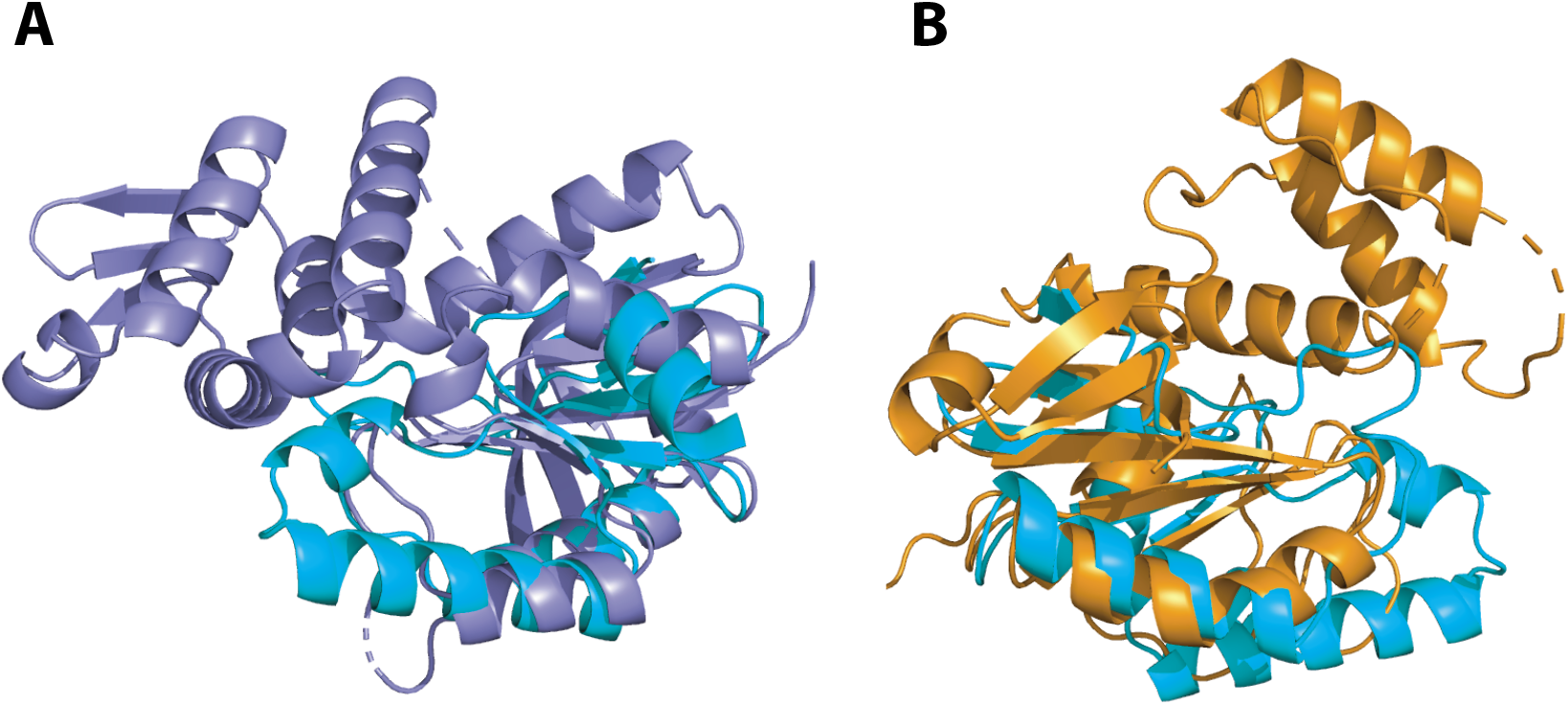
Superimposition of the nuclease-like domain of Can1 (cyan) with the nuclease domain of (A) VC1899 (mauve) and (B) Can1 (orange). The nuclease-like domain superimposes with around half of the secondary structure elements found in the nuclease domain. It lacks the N-terminal α-helix, β-strand, three α-helices (in a helix-loop-helix-loop-helix arrangement) and β-strand. The nuclease-like domain overlaps well with the next two β-strands, α-helix, and β-strand, and then has an insertion (with respect to the nuclease) of a helix-turn-helix motif followed by a β-strand. The final α-helix of both domains superimposes well.

## References

1. Rouillon, C., Athukoralage, J.S., Graham, S., Grüschow, S. & White, M.F. Control of cyclic oligoadenylate synthesis in a type III CRISPR system. eLife 7, e36734 (2018).

2. Niewoehner, O. et al. Type III CRISPR-Cas systems produce cyclic oligoadenylate second messengers. Nature 548, 543–548 (2017).

3. Kazlauskiene, M., Kostiuk, G., Venclovas, C., Tamulaitis, G. & Siksnys, V. A cyclic oligonucleotide signaling pathway in type III CRISPR-Cas systems. Science 357, 605–609 (2017).

4. Nasef, M. et al. Regulation of cyclic oligoadenylate synthesis by the S. epidermidis Cas10-Csm complex. RNA, pii: rna.070417.119 (2019).

5. Koonin, E.V. & Makarova, K.S. Discovery of Oligonucleotide Signaling Mediated by CRISPR-Associated Polymerases Solves Two Puzzles but Leaves an Enigma. ACS Chem Biol 13, 309–312 (2018).

6. Niewoehner, O. & Jinek, M. Structural basis for the endoribonuclease activity of the type III-A CRISPR-associated protein Csm6. RNA 22, 318–29 (2016).

7. Foster, K., Kalter, J., Woodside, W., Terns, R.M. & Terns, M.P. The ribonuclease activity of Csm6 is required for anti-plasmid immunity by Type III-A CRISPR-Cas systems. RNA Biol, 1–12 (2018).

8. Sheppard, N.F., Glover, C.V., 3rd, Terns, R.M. & Terns, M.P. The CRISPR-associated Csx1 protein of Pyrococcus furiosus is an adenosine-specific endoribonuclease. RNA 22, 216–24 (2016).

9. Han, W., Pan, S., Lopez-Mendez, B., Montoya, G. & She, Q. Allosteric regulation of Csx1, a type IIIB-associated CARF domain ribonuclease by RNAs carrying a tetraadenylate tail. Nucleic Acids Res 45, 10740–10750 (2017).

10. Athukoralage, J.S., Graham, S., Grüschow, S., Rouillon, C. & White, M.F. A type III CRISPR ancillary ribonuclease degrades its cyclic oligoadenylate activator. J. Mol. Biol. in press(2019).

11. Lintner, N.G. et al. The structure of the CRISPR-associated protein Csa3 provides insight into the regulation of the CRISPR/Cas system. J Mol Biol 405, 939–55 (2011).

12. Athukoralage, J.S., Rouillon, C., Graham, S., Grüschow, S. & White, M.F. Ring nucleases deactivate Type III CRISPR ribonucleases by degrading cyclic oligoadenylate. Nature 562, 277–280 (2018).

13. Makarova, K.S., Anantharaman, V., Grishin, N.V., Koonin, E.V. & Aravind, L. CARF and WYL domains: ligand-binding regulators of prokaryotic defense systems. Front Genet 5, 102 (2014).

14. Deng, L., Garrett, R.A., Shah, S.A., Peng, X. & She, Q. A novel interference mechanism by a type IIIB CRISPR-Cmr module in Sulfolobus. Mol Microbiol 87, 1088–99 (2013).

15. Hatoum-Aslan, A., Maniv, I., Samai, P. & Marraffini, L.A. Genetic Characterization of Antiplasmid Immunity through a Type III-A CRISPR-Cas System. Journal of Bacteriology 196, 310–317 (2014).

16. Jiang, W., Samai, P. & Marraffini, L.A. Degradation of Phage Transcripts by CRISPR-Associated RNases Enables Type III CRISPR-Cas Immunity. Cell 164, 710–21 (2016).

17. Rostol, J.T. & Marraffini, L.A. Non-specific degradation of transcripts promotes plasmid clearance during type III-A CRISPR-Cas immunity. Nat Microbiol 4, 656–662 (2019).

18. Staals, R.H. et al. RNA targeting by the type III-A CRISPR-Cas Csm complex of Thermus thermophilus. Mol Cell 56, 518–30 (2014).

19. Staals, R.H. et al. Structure and activity of the RNA-targeting Type III-B CRISPR-Cas complex of Thermus thermophilus. Mol Cell 52, 135–45 (2013).

20. Taylor, D.W. et al. Structural biology. Structures of the CRISPR-Cmr complex reveal mode of RNA target positioning. Science 348, 581–5 (2015).

21. Liu, T.Y., Iavarone, A.T. & Doudna, J.A. RNA and DNA Targeting by a Reconstituted Thermus thermophilus Type III-A CRISPR-Cas System. PLoS One 12, e0170552 (2017).

22. Shah, S.A. et al. Comprehensive search for accessory proteins encoded with archaeal and bacterial type III CRISPR-cas gene cassettes reveals 39 new cas gene families. RNA Biol, 1–13 (2018).

23. Agari, Y. et al. Transcription profile of Thermus thermophilus CRISPR systems after phage infection. J Mol Biol 395, 270–81 (2010).

24. Zimmermann, L. et al. A Completely Reimplemented MPI Bioinformatics Toolkit with a New HHpred Server at its Core. J Mol Biol 430, 2237–2243 (2018).

25. Rouillon, C., Athukoralage, J.S., Graham, S., Grüschow, S. & White, M.F. Investigation of the cyclic oligoadenylate signalling pathway of type III CRISPR systems. Methods Enzymol 616, 191–218 (2019).

26. Yang, W. Nucleases: diversity of structure, function and mechanism. Q Rev Biophys 44, 1–93 (2011).

27. Long, F. et al. AceDRG: a stereochemical description generator for ligands. Acta Crystallogr D Struct Biol 73, 112–122 (2017).

28. Holm, L. & Laakso, L.M. Dali server update. Nucleic Acids Res 44, W351–5 (2016).

29. Tamulaitiene, G. et al. Restriction endonuclease AgeI is a monomer which dimerizes to cleave DNA. Nucleic Acids Res 45, 3547–3558 (2017).

30. Deibert, M., Grazulis, S., Sasnauskas, G., Siksnys, V. & Huber, R. Structure of the tetrameric restriction endonuclease NgoMIV in complex with cleaved DNA. Nat Struct Biol 7, 792–9 (2000).

31. Varble, A. & Marraffini, L.A. Three New Cs for CRISPR: Collateral, Communicate, Cooperate. Trends Genet 35, 446–456 (2019).

32. Abudayyeh, O.O. et al. C2c2 is a single-component programmable RNA-guided RNA-targeting CRISPR effector. Science 353, aaf5573 (2016).

33. Meeske, A.J. & Marraffini, L.A. RNA Guide Complementarity Prevents Self-Targeting in Type VI CRISPR Systems. Mol Cell 71, 791–801 e3 (2018).

34. Meeske, A.J., Nakandakari-Higa, S. & Marraffini, L.A. Cas13-induced cellular dormancy prevents the rise of CRISPR-resistant bacteriophage. Nature (2019).

35. Elmore, J.R. et al. Bipartite recognition of target RNAs activates DNA cleavage by the Type III-B CRISPR-Cas system. Genes Dev 30, 447–59 (2016).

36. Estrella, M.A., Kuo, F.T. & Bailey, S. RNA-activated DNA cleavage by the Type III-B CRISPR-Cas effector complex. Genes Dev 30, 460–70 (2016).

37. Kazlauskiene, M., Tamulaitis, G., Kostiuk, G., Venclovas, C. & Siksnys, V. Spatiotemporal Control of Type III-A CRISPR-Cas Immunity: Coupling DNA Degradation with the Target RNA Recognition. Mol Cell 62, 295–306 (2016).

38. Samai, P. et al. Co-transcriptional DNA and RNA Cleavage during Type III CRISPR-Cas Immunity. Cell 161, 1164–74 (2015).

39. Chen, J.S. et al. CRISPR-Cas12a target binding unleashes indiscriminate single-stranded DNase activity. Science 360, 436–439 (2018).

40. Li, S.Y. et al. CRISPR-Cas12a has both cis- and trans-cleavage activities on singlestranded DNA. Cell Res 28, 491–493 (2018).

41. Linkert, M. et al. Metadata matters: access to image data in the real world. J Cell Biol 189, 777–82 (2010).

42. Schindelin, J. et al. Fiji: an open-source platform for biological-image analysis. Nat Methods 9, 676–82 (2012).

43. Winter, G. xia2: an expert system for macromolecular crystallography data reduction. Journal of Applied Crystallography 43, 186–190 (2010).

44. Kabsch, W. Xds. Acta crystallographica. Section D, Biological crystallography 66, 125–32 (2010).

45. Adams, P.D. et al. PHENIX: a comprehensive Python-based system for macromolecular structure solution. Acta crystallographica. Section D, Biological crystallography 66, 213–21 (2010).

46. Murshudov, G.N., Vagin, A.A. & Dodson, E.J. Refinement of macromolecular structures by the maximum-likelihood method. Acta Crystallogr D Biol Crystallogr 53, 240–55 (1997).

47. Winn, M.D. et al. Overview of the CCP4 suite and current developments. Acta Crystallogr D Biol Crystallogr 67, 235–42 (2011).

48. Emsley, P., Lohkamp, B., Scott, W.G. & Cowtan, K. Features and development of Coot. Acta Crystallogr D Biol Crystallogr 66, 486–501 (2010).

49. Chen, V.B. et al. MolProbity: all-atom structure validation for macromolecular crystallography. Acta crystallographica. Section D, Biological crystallography 66, 12–21 (2010).

50. Basham, M. et al. Data Analysis WorkbeNch (DAWN). J Synchrotron Radiat 22, 853–8 (2015).

51. Brunger, A.T. Version 1.2 of the Crystallography and NMR system. Nat Protoc 2, 2728–33 (2007).

52. Schneidman-Duhovny, D., Hammel, M., Tainer, J.A. & Sali, A. FoXS, FoXSDock and MultiFoXS: Single-state and multi-state structural modeling of proteins and their complexes based on SAXS profiles. Nucleic Acids Res 44, W424–9 (2016).

53. Schneidman-Duhovny, D., Hammel, M., Tainer, J.A. & Sali, A. Accurate SAXS profile computation and its assessment by contrast variation experiments. Biophys J 105, 962–74 (2013).

